# A Comparative Study of Muscular, Vestibular, and Haptic Stimulation on Dream Incorporation

**DOI:** 10.1101/2024.12.05.626974

**Authors:** Emma Peters, Jakob Pöhlmann, Xinlin Wang, Martin Dresler, Daniel Erlacher

## Abstract

The connection between the dreamed body and the real physical body remains a subject of ongoing investigation. This study explored how the dreamed body responds to somatosensory stimulation of the physical body, aiming to shed light on the sensory processes that shape our dreaming experiences. We employed a novel within-subject design to compare the incorporation of three different types of bodily stimuli—electrical muscular, galvanic vestibular, and haptic vibration—into dream content, alongside a control sham condition for each stimulus. In total, 24 participants spent one adaptation night, followed by three consecutive test nights in the sleep laboratory. REM awakenings, after sham or stimulation periods, were carried out for dream report collection. In total, 165 dream reports were collected across conditions. While dream incorporation was observed across the three stimulation methods, it occurred equally in both the stimulation and sham conditions for all three modalities. These findings highlight broader methodological challenges in dream incorporation research and raise concerns about potential confounding factors affecting the interpretation of results. Future research with larger sample sizes is needed to detect smaller effect sizes and fully understand the influence of these somatosensory stimuli on dream content. This study employed a rigorous experimental approach to exploring dream incorporation and addressed many methodological challenges in this area. We further suggest areas of improvement to optimize dream incorporation of different sensory modalities.

**Graphical Abstract:** 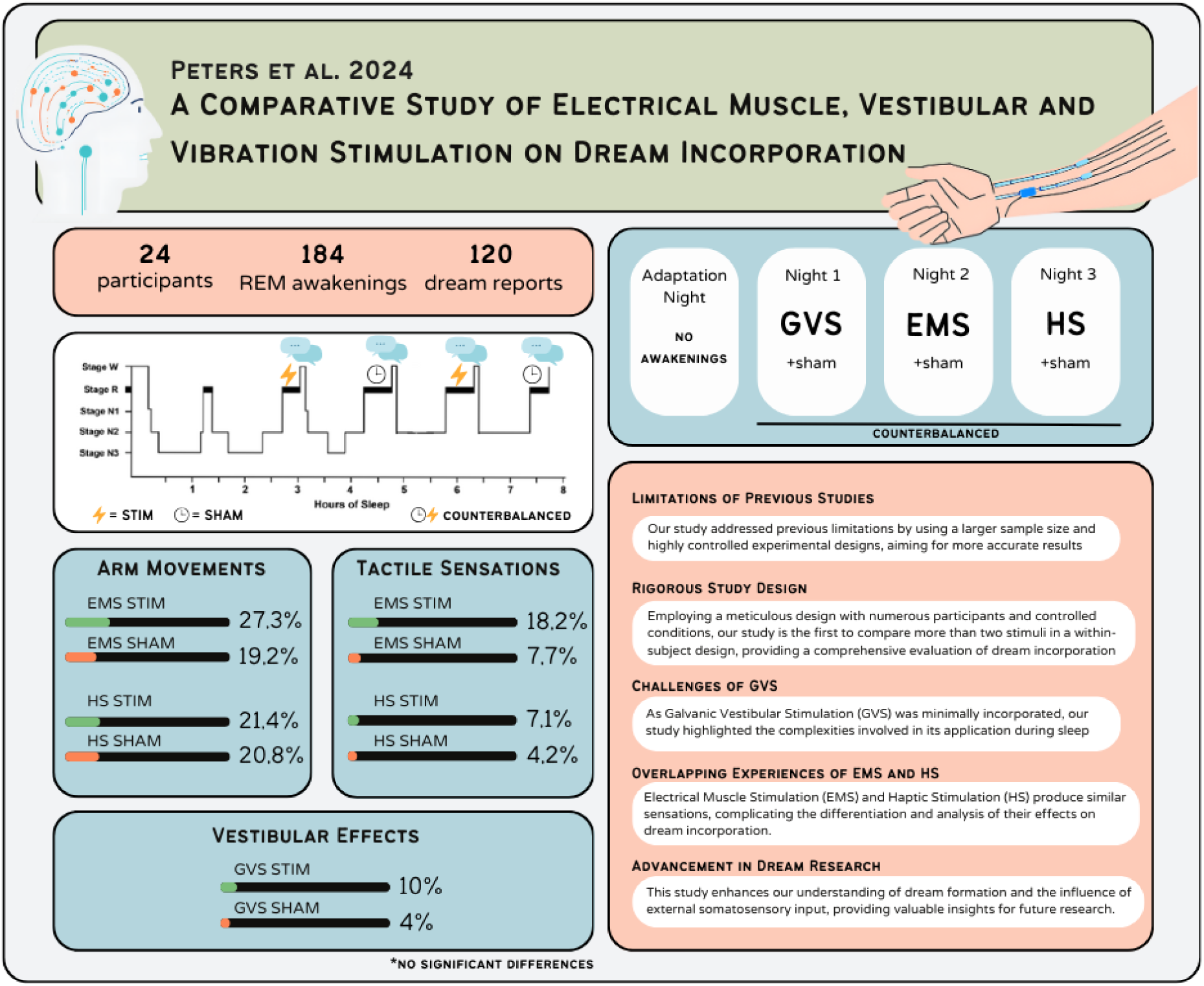

## Introduction

Experimental alteration of dream content using external stimulation during sleep has been explored for many decades as dream content formation is influenced by external environmental stimuli and internal bodily cues. They have the potential to infiltrate our dreamscapes, shaping the content and experiences we encounter during sleep. This phenomenon, known as dream incorporation, demonstrates the brain’s ability to integrate sensory information such as sounds, scents, light, or even temperature, into the dream (Salvesen et al., 2024; Solomonova & Carr, 2019). Understanding the impact of external stimuli on dreams is crucial for uncovering the fundamentals of dream formation and the role of the physical body in shaping these experiences during sleep. Additionally, dream engineering approaches might enable the alteration of dream content for clinical purposes, allowing interventions in dream content to address various therapeutic needs (Vitali et al., 2022). Modifying dream content through external stimulation can serve as an initial step toward inducing lucid dreaming, a fascinating phenomenon where the dreamer becomes aware of dreaming while still asleep (Dresler et al., 2016; Zerr et al., 2024). By accessing the dreamer, external stimulation can act as a lucidity cue, prompting the reflection necessary for dream awareness (Tan & Fan, 2022). Lucid dreaming has applications across various fields, including nightmare therapy, creative problem solving, skill development and personal exploration (Ouchene et al., 2023; Peters et al., 2023; Schädlich & Erlacher, 2012).

### The Role of the Sensorimotor Cortex and Bodily Synchrony in Dream Experiences

Recent advances in neuroscience have shed light on the neurophysiological mechanisms underlying the interconnectedness of the dreamed and real body. Research has demonstrated that specific dream actions, such as hand clenching, are associated with activation of the sensorimotor cortex, mirroring patterns observed during wakeful behaviour (Dresler et al., 2011). Later, the mechanisms driving these dream contents were investigated by using transcranial direct current stimulation (tDCS). The effects of transcranial direct current stimulation (tDCS) on dream content vary depending on stimulation location and parameters. Earlier studies examined the effects of tDCS on motor imagery during the waking resting state. Anodal unihemispheric tDCS over the motor cortex (C3) increased general and athletic motor imagery compared to sham stimulation, demonstrating tDCS’s ability to modulate motor imagery without active motor tasks, with potential applications in performance enhancement and rehabilitation (Speth et al., 2015). Extending this research to REM sleep, anodal unihemispheric tDCS of the left motor cortex (C3) was found to enhance the quantity and quality of athletic motor imagery during dreams compared to cathodal and sham stimulation, suggesting that REM sleep motor imagery might function as rehearsal for motor movements (Speth & Speth, 2016). Conversely, bihemispheric tDCS targeting the primary motor cortex during REM sleep significantly reduced reports of dream movements, particularly repetitive actions, without altering tactile or vestibular sensations, implicating the sensorimotor cortex in generating movement sensations in dreams (Noreika et al., 2020). Besides neural activations, physiological synchrony between dreamed actions and the physical body has been observed. Research suggests that dreamed actions involving eye gaze and facial expressions exhibit physiological parallels with the physical body (Arnulf, 2011; Dement & Kleitman, 1957). Similarly, facial muscle activations associated with emotions, such as smiling or frowning, have been observed to mirror the dream content (Rivera et al., 2018). During rapid eye movement (REM) sleep, which is associated with vivid dreaming, bodily responses such as changes in heart rate, respiration, and muscle activity can mirror the experienced dream content (Erlacher & Schredl, 2008). Dreamed actions thus influence the physical, sleeping body, and it seems this is bidirectional. The synchronization implies an interactive process where external stimuli affecting the physical body—like loud noises or touch—can influence dream experiences, linking the dream body with the sleeping person’s physical state. Thus, the dreamed body appears indirectly connected to the sleeper’s surroundings and experiences. Overall, these neural and physiological links highlight the continuity between wakefulness and dreaming, revealing the sleeping brain’s active role in integrating external stimuli into dreams.

### Influence of Somatosensory Stimulation on Dream Content Incorporation

A wide range of studies have been done, testing different external stimuli and their success in incorporating them into one’s dream. The stimuli ranged from somatosensory to visual, auditory, and olfactory applications (Salvesen et al., 2024). Dream incorporation often involves exploring distant sensory stimulation, which refers to sensory inputs that do not physically touch the body but are perceived and experienced during dreams, as opposed to proximal sensory stimulation, stimuli that directly contact the dreamer’s body. Recognizing the strong interconnection between the dreamed body and the physical body led researchers to show interest in exploring the effects of somatosensory stimuli.

The impact of electrical muscle stimulation on dreams by administering electrical impulses to participants’ wrists during sleep was previously investigated. The study included 10 male participants, evenly split between night sleepers and night-shift workers accustomed to daytime sleep. The experimental setup followed participants’ natural sleep schedules in a laboratory setting. A variety of different stimulation conditions were employed: stimulation immediately upon detection of REM sleep (C2, C3) or 3 minutes afterward (C4, C5), with awakenings scheduled either 30 seconds (C2, C4) or 3 minutes after stimulation (C3, C5). A control sham condition (C1) involved awakening 3 minutes after REM onset. Findings indicated that early REM stimulation was most effective in altering dream experiences, while later REM stimulation increased the likelihood of waking. Dreams from condition C3 exhibited significantly higher ratings for ‘body centrality’ and ‘body activity,’ with C4 also showing elevated ‘body activity’ ratings. Direct and indirect incorporation rates combined in conditions C2, C3, C4, and C5 were notably higher compared to the control condition (C1) (Koulack, 1969). Tactile somatosensory stimulation has been investigated using vibrotactile stimuli on the index finger (N=10), and wrist or ankle (N=14). Each stimulus was repeated up to five times after at least 5 minutes of confirmed REM sleep, starting from the third REM cycle of the night. Participants were awakened by the experimenter 1 minute after the fifth stimulus. Results indicated that 9 out of 21 dream reports following index finger stimulation (42.9%) and 13 out of 27 dream reports following wrist and ankle stimulation (48.1%) incorporated the stimulus, as reported by the participants. This study however did not employ any control conditions (Paul et al., 2014).

The involvement of the vestibular system in dream incorporation has been explored by using a hammock and a tilted bed. Leslie and Ogilvie (1996) had seven participants sleep in a hammock, which rocked 30° from vertical at 1 Hz, starting 5 minutes into the second or later REM sleep cycles on one of two experimental nights—the other night serving as a control. Participants were awakened after 5 minutes of stimulation or stationary control, resulting in 45 dream reports. Participants were asked about stimulus perception and the dream reports were scored by external raters on the presence of vestibular experiences involving the body, kinaesthetic feelings involving the vestibular system, bizarre trajectories of other objects, psychological feelings of disorientation and other possible effects of regular rocking. A higher rate of stimulus incorporation was noted for the rocking condition (25%) compared to the control condition (7%). Additionally, there was a significant correlation between vestibular incorporation and dream bizarreness ratings (Leslie & Ogilvie, 1996). The impact of tilting the upper part of a bed by 10° on sleep and dreams in two nap sessions was also studied, involving the bed being tilted versus having the bed in a horizontal position as the control condition. The bed was tilted 20 minutes after sleep onset, and participants were awakened shortly after. Upon awakening, participants categorized their experiences as ‘usual thought,’ ‘hypnagogic imagery,’ ‘no experience,’ ‘uncertain/forgotten,’ ‘dream,’ or ‘sleep paralysis.’ External raters found no differences in the amount of vestibular or somatosensory information between conditions (Nozoe et al., 2020) No further detailed analysis of dream content was conducted, which could have provided valuable insights.

On average, the incorporation rates for all stimuli incorporation across studies seem to be ranging from 0% to 90% (Salvesen et al., 2024; Solomonova & Carr, 2019), with tactile and auditory stimulations being the most successful ones, however some of the studies lack methodological rigor and the results should be approaches with caution.

### Research question and hypotheses

Decades of research have investigated dream incorporation, however the results relevant for the current study were either based on a low number of participants or lacked a thorough control condition. In addition, very few have attempted comparisons of more than two stimuli and looked at their efficiency and specificity in altering dream content. In this study, somatosensory stimulation will be explored using three different modalities: muscle, haptic, and vestibular stimuli, in a within-subject design including a control condition for each modality, a novel, rigorous, and challenging approach. We aim to use the physical, sleeping body as a means to access the dreamed body and alter the dream experience using these approaches. The three stimulation methods are galvanic vestibular stimulation (GVS), electrical muscle stimulation (EMS), and haptic vibratory stimulation (HS). We hypothesize that the three stimuli will be incorporated into the dream, targeting their respective modalities. We expect the galvanic stimulation to increase vestibular and balance imagery in dreams (e.g., flying or falling); muscle stimulation to increase bodily movement in the dream (e.g., limb movement, whole-body movement); and haptic stimulation to increase somatosensory effects (e.g. pressure, vibration, etc.) This approach allows for robust comparisons and a thorough examination of how different sensory stimuli influence dream experiences, aiming to provide a more comprehensive understanding of the interplay between external stimulation and dream content integration while approaching dream incorporation methodology through a critical lens.

## Methods

### Participants

Participants (n=24; 12 female) were recruited through advertisements around the campus of Bern University and via social media groups and circles of the researchers around Bern, Switzerland. Participants were healthy and between 20 - 37 years old (M = 26.5, SD = 4.97) and able to either speak English, German, or Swiss-German. When applying for the study, participants were pre-screened online and excluded if they had a low dream recall rate (<3 per week), late bedtime (>1 am), long sleep latency (>1 hour), and finally, if they, or any family members, were diagnosed with epilepsy, sleep-related disorders or current psychiatric disorders. The average dream recall was 4 days per week. An a priori calculation was performed to calculate the optimal sample size, resulting in an effect size of w = 0.35, alpha = .05, power = 0.8, and df = 2 (three conditions), revealing an optimal sample size of 26 using a within-subject design.

### Measures and Procedures

The study was performed in the sleep laboratory at the Institute of Sport Sciences (University of Bern). Participants were financially compensated. The Faculty of Human Sciences of the University of Bern approved the study ethically. Before the study, participants were informed about the aim of the study and signed a written consent form. Data protection was ensured by using a pseudonymous code for each participant.

### Experimental Setup

Each participant spent four consecutive nights in the sleep laboratory. The participant arrived at 9 pm each night and got ready for bed. Polysomnography (PSG) was administered to monitor sleep stages, consisting of electroencephalography (EEG), electrooculography (EOG), and electromyography (EMG). PSG set-up was applied according to the 10-20 system (Klem et al., 1999). F3, F4, C3, C4, O1, O2, horizontal EOG1 and EOG2, chin EMGl, chin EMGr, a reference (earlobe) and ground (Cz) electrodes were used. The first night was an adaptation night, with the function of getting the participants used to the environment and the PSG set-up. During this first night, no stimulation devices were applied, and no awakenings were performed. In each of the following three nights, participants received one of the three stimuli (haptic, vestibular, muscle) during REM sleep periods, including sham stimulation trials. One stimulation device was applied per night, and the order of the stimulation methods and sham stimulation was randomized and counterbalanced, as can be seen in Figure 1.

**Figure 1:**
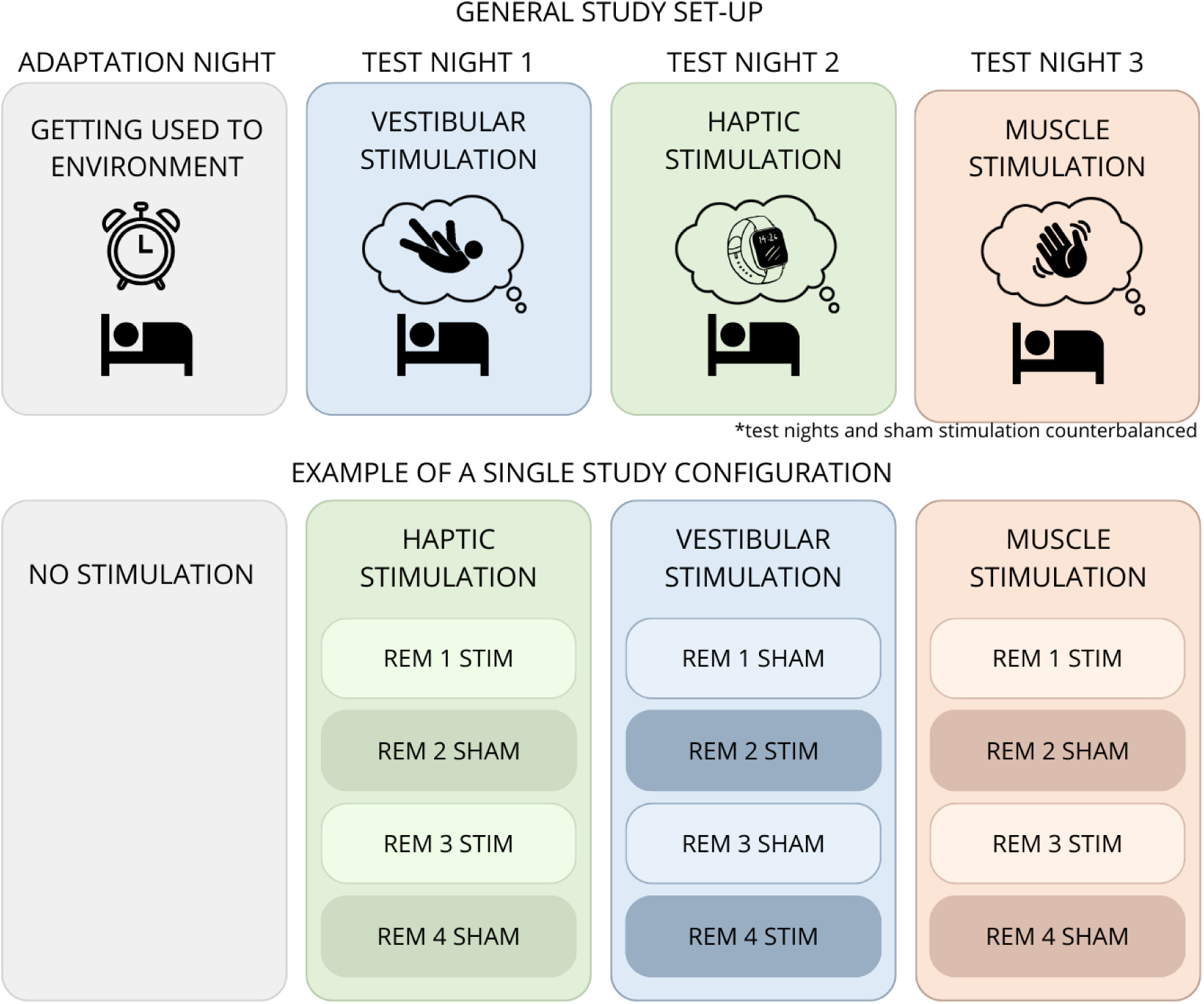
Top: the study overview with the adaptation night and the three stimulation nights. Bottom: an example of the configuration for a single participant with STIM and SHAM order in REM phases balanced across the nights. In this specific configuration, the night order is haptic, vestibular and muscle stimulation.

### Stimulation Methods

The three different stimuli that were used consisted of Galvanic Vestibular Stimulation (GVS), Electrical Muscle Stimulation (EMS) and Haptic Stimulation (HS). Each stimulation lasted for two seconds.

*(Sinusoidal) Galvanic Vestibular Stimulation* was set up using the NeuroConn DC-Stimulator Plus (NeuroConn DC-STIMULATOR PLUS | Neurocare Technology, 2023) and by attaching two carbon rubber electrodes behind the ear over the mastoid bone. Electrical Stimulation over the mastoid bone stimulates the otoliths, which are part of the vestibular system (Khosravi-Hashemi et al., 2019; Sra et al., 2019). This can induce a variety of vestibular sensations, depending on the position of the body and head in space, as well as the electrode placement (Khosravi-Hashemi et al., 2019). In order to find a consistent stimulation experience across participants, we investigated GVS in detail. The effect of the stimulation on horizontal body position was tested, including positions, on the back, on the side, using a pillow or no pillow. In addition to spatial positioning, different stimulation parameters were tested, including stimulation intensity (μA), duration (s), frequency (Hz) and type of waveform (tDCS, tACS, sines, half-sines), fade-in and fade-out time (s). We noticed that the frequency of the stimulation had the most consistent effect on the perception, in which lower frequencies (0.5Hz) resulted in a boat or wave-like feeling, while higher frequencies (1-2Hz) were perceived as a more abrupt, unnatural, and externally produced movement. The final intensity was set to a level where the participants felt a small sway sensation without feeling an uncomfortable skin sensation from the current behind the ear while sitting on the bed with their eyes open. The stimulus was set to a frequency of 1 Hz and lasted for two seconds, resulting in a quick left-right-left-right sway sensation of the head.

*Electrical Muscle Stimulation* induced a movement of the wrist and fingers consisting of a flexion of the ring and pinky finger. To achieve this, two self-adhesive electrodes were placed on the left? forearm of the participant, near the origin of the wrist flexor muscles. The device used to apply the electrical stimulation was the Rehamove 3 “ScienceMode” Device. The Rehamove 3 is a small handheld device that can be connected to a computer, enabling the stimulation trigger externally (Kubersky, 2018; *RehaMove—Funktionelle Elektrostimulation | HASOMED GmbH.*, 2023). The stimulus was tested before going to bed while the participant was still awake to establish the lowest possible intensity that induced flexion of the ring and pinky finger, as well as to ensure that the stimulation was comfortable for the participant. This intensity was kept throughout the night.

*Haptic Stimulation* was applied using a home-made device. This consisted of a small plastic tube with a rotational vibrational motor inside connected to a current-producing Lab Jack U3 LV that could be triggered from the control room to produce a light vibration of the plastic tube. The vibrating device was taped to the left dorsal wrist and the cable was further secured by taping it to the arm. The stimulus was tested prior to bedtime to familiarize the participants with the sensation, comparable to that of a vibrating phone. The stimulation only had one intensity setting in contrast to EMS.

### Stimulation Protocol

The stimulation was carried out during REM sleep, using one of the randomly selected stimulation methods in a counterbalanced order. The first REM phase was disregarded, and the REM awakenings were carried out from the second REM phase onwards. Depending on whether it was the second, third, or fourth REM phase, the researcher had to wait five, ten, or 15 minutes respectively, ensuring REM stability and thus likely formation of a dream plot. Either stimulation (STIM) or no stimulation (SHAM) was performed, again, in a counterbalanced order. All stimulations were triggered from the control room, ensuring minimal disturbance for the participant. Every STIM condition contained a maximum of 10 two-second stimulations, with a 30-second wait period in between each stimulation. After the final stimulation, there was another wait period of 30 seconds before the participant was awakened. The SHAM condition consisted of a timer that was set to match the duration of the entire STIM condition. If the participant went into NREM, the timer was aborted and started again at the following REM onset. If the participant woke up completely due to arousal following stimulation or during the SHAM timer, the protocol was aborted, and the dream report was collected. When the participant stayed in REM sleep throughout the 10 stimulations, the researcher awakened the participant via intercom, and a dream report was collected. Answers were recorded on audiotape and later transcribed. After collecting the dream report, the participants were allowed to go back to sleep, the researcher waited for the next REM phase, and repeated the protocol.

### REM Awakenings

The REM awakenings started by calling the participant by their name, followed by asking if they are awake. When they reported being awake, the question: ‘What went through your head before you woke up?’ was asked. After the participant finished their report, the question: ‘Did you notice anything else?’ was asked in order to collect more detail. Then, participants were asked directly about the perception of the stimulation of the dream. For EMS and HS; the question was: ‘Did you notice the stimulation in the dream?’, and for GVS: ‘Did you notice unusual balance in the dream?’. Finally, the question: ‘Were you aware that you were dreaming while you were dreaming?’ was asked with the aim of assessing the presence of any lucid dreams.

### Dream Data Analysis

After transcribing the audio dream reports to written form, they were processed following a dream report cleaning Manual (Schredl, 2010). These include the removal of any words or sentences that do not directly refer to the dream content. Because the dream reports were collected in multiple different languages, they were all translated into English using DeepL Translator to ensure high-quality and contextually appropriate translations. Finally, the dream reports were randomized and rated by an external, blind rater. To evaluate the dream reports, the rater used a Dream Content Analysis (DCA) Manual, which contained stimuli-related incorporation of movement, balance, and somatosensory sensations. In addition to these three core scales, scales on realism/bizarreness (Schredl, 2003), the number of dream characters, interaction, emotions (Schredl, 2018), and the incorporation of the environment (sleep laboratory, electronic devices, experiment, researcher) taken from the Bodily Experiences in Dreams (BED) Questionnaire (Noreika et al., 2020) were included to assess other dream attributes. Direct incorporation of laboratory content was investigated using the scale: ‘Are there any references to the sleep laboratory environment, electrodes, researcher, etc. present in the dream?’, answered by a ‘yes’ or ‘no.’ Each dream report was evaluated quantitatively depending on these various categories. Any dreams reporting the participant being awake during stimulation or waking up as a result of stimulation were excluded.

For each of the three stimulation methods, one scale was selected as representing direct incorporation for that stimulus. For GVS, the scale: ‘General Balance-Related Activities’ was selected. This scale is phrased in the Dream Content Analysis Manual as: ‘Does the dreamer perform activities involving balancing and balance, e.g. flying, going over a narrow bridge? Does the dreamer have problems with balance, e.g. the occurrence of dizziness, fear of looking deep, etc?’. The scale thus combines all these parameters. For EMS: ‘Movement of the Arms’ was selected and was phrased as: ‘Does the dreamer perform activities that have to do with the arms, e.g., B., waving, manual labour, boxing? The activity should be explicitly named.’ Finally, for HS, the first scale: ‘Does the dreamer experience activities in which a tactile or somatosensory sensation/stimulus plays a role?’ was selected together with a second scale: ‘Mark the parts of the body where the sensation occurred in the first scale.’ If the first scale was scored as a 1 in combination with the mention of the arms in the second scale, then the final score for ‘Tactile or Somatosensory Sensations in the Arms.’ was scored as 1. Dream incorporation rate was calculated by taking the sum of all scores per stimulation category and correcting them for the number of dreams per category.

### Polysomnography Data

To objectively verify the presence of REM sleep during the stimulation trials, we exported and assessed the PSG data. Dream reports were excluded if wakefulness was detected during the stimulation trials. Instances where stimulation failed due to technical issues were classified as SHAM trials.

### Statistics

The dream incorporation rate for the three stimulation methods in the relevant scales was compared using a mixed model statistical test using RStudio 2023.6.1, fitting all models with the package lme4 (Bates et al., 2015). ANOVA tests were performed to test each linear mixed-effects model for significance, and post hoc pairwise comparisons were conducted using the ‘*emmeans*’ (version 1.8.7) function in the RStudio package with Tukey correction. We applied separate mixed models on the scales: ‘Movement of the Arms,’ ‘Tactile or Somatosensory Sensations in the Arm’, and ‘General Balance-Related Activities.’ The conditions (EMS, GVS, HS) and stimulation (SHAM or STIM) were used for fixed factors, A by-subject random intercept was added to represent between-person variability and account for the unbalanced data structure (Bates et al., 2015). To further explore whether the current sample size could efficiently detect the significant effects on these models, we performed power analysis on mixed models on each scale.

## Results

### Awakenings and Dream reports

In 72 experimental nights, excluding adaptation nights, a total of 184 awakenings were performed in REM sleep, 96 in the STIM condition, and 88 in the SHAM condition. 65, 61, and 58 awakenings were performed in the EMS, GVS, and HS conditions, respectively. A total of 165 dream reports were recorded, which is equivalent to a dream recall rate of 89.7%. Forty-five dream reports were excluded due to awakening during stimulation based on the dream reports and the PSG data; the remaining 120 dream reports were included in the analysis. 45 Dreams were included in the STIM condition and 75 in the SHAM condition. One participant did not report any dreams throughout the experiment. The average dream length of the EMS STIM group was 59.2 words, for EMS SHAM, this was 68.7. For GVS STIM and GVS SHAM this was 91.9 and 88.1 words respectively. Finally, HS STIM and HS SHAM had an average dream length of 72.7 and 79.7 words respectively. An ANOVA-test revealed that none of these results differed significantly.

### Dream Incorporation

Participants were asked directly about the perception of the stimulation of the dream. In the EMS STIM condition, 18.2% of dreams contained perception of the stimulation in the dream, followed by 3.8% in the EMS SHAM condition. In the GVS STIM and GVS SHAM conditions, 20% and 4% of dreams contained a distinct vestibular experience. Finally, In the HS STIM condition, 21.4% of dreams contained perception of the stimulation, whereas, in the HS SHAM condition, this was 12.5% of dreams. A Fischer test proved that none of these percentages were significantly different between conditions. Using the DCA manual, a variety of scales were investigated. The DCA manual and the results can be found in the Appendix. For our research question, we will consequently focus on three main scales representing the relevant stimulation method. For EMS and HS, this was ‘Movement of the Arms’ and ‘Tactile or Somatosensory Sensations in the Arm’, and for GVS, this was ‘General Balance-Related Activities’. EMS and HS are tested on the same scales as they have overlapping stimulation experiences, whereas GVS stimulation elicited a completely independent experience. Figure 2 presents the dream incorporation rate (DIR), which is the total dream incorporation, corrected for the number of dreams collected per condition.

**Figure 2:**
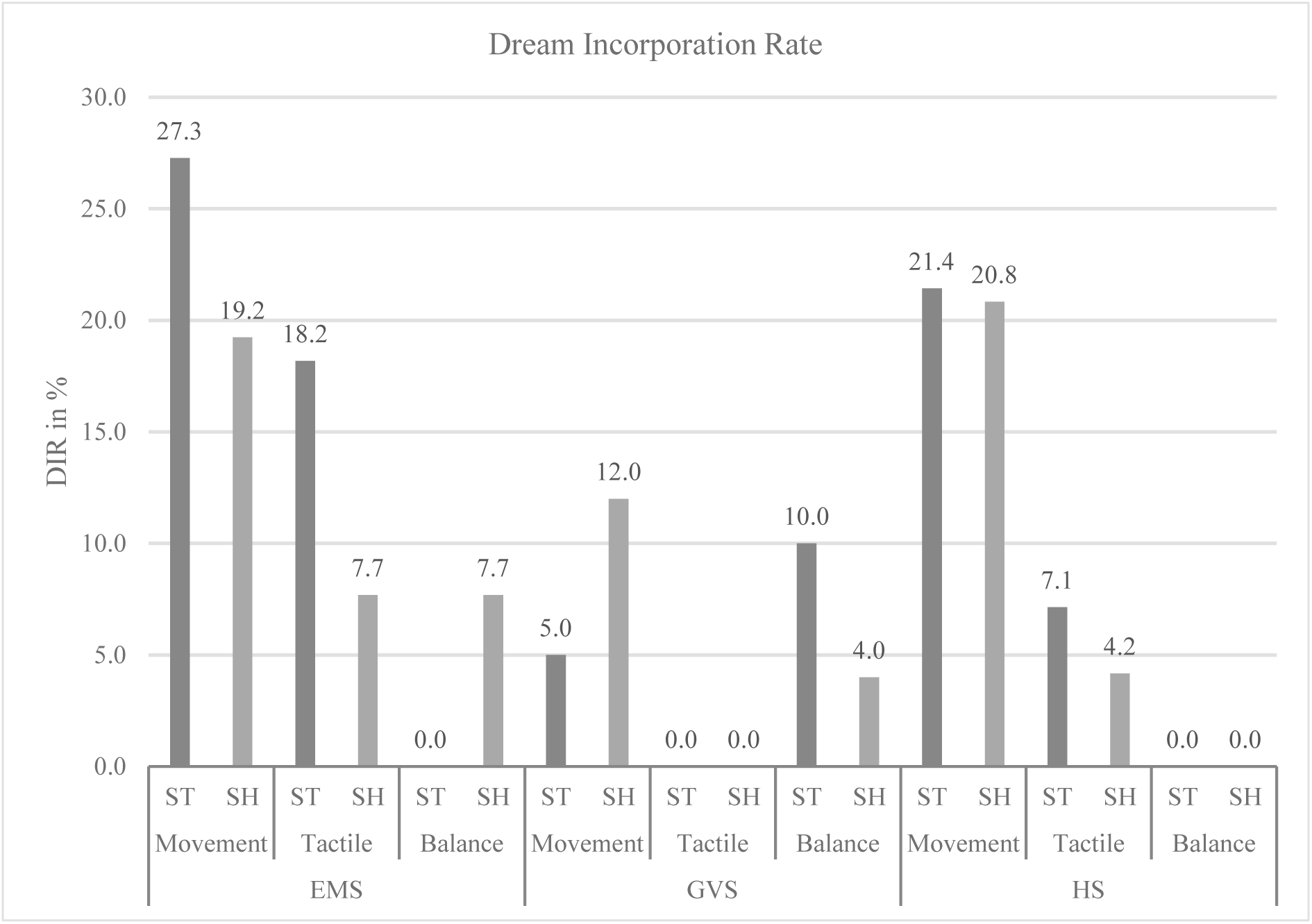
The Dream Incorporation Rate (DIR) of ‘Movement of the Arms,’ ‘Tactile or Somatosensory Sensations in the Arms,’ and ‘General Balance-Related Activities’ per stimulation condition. ST = stimulation condition, SH = sham condition. EMS: Electrical Muscle Stimulation, GVS: Galvanic Vestibular Stimulation, HS: Haptic Stimulation, STIM = stimulation condition, SHAM: sham stimulation condition.

The dream incorporation rates of different stimuli were evaluated across six conditions: EMS STIM, EMS SHAM, GVS STIM, GVS SHAM, HS STIM, and HS SHAM. The results indicate varying levels of incorporation of ‘Movement of the Arms,’ ‘Tactile or Somatosensory Sensations in the Arms,’ and ‘General Balance-Related Activities.’ In the EMS STIM condition, 27.3% of dreams incorporated arm movements, 18.2% included Tactile or Somatosensory Sensations in the Arms, and no dreams included General Balance-Related Activities. The EMS SHAM condition showed 19.2% incorporation for arm movements, 7.7% for tactile sensations, and 7.7% for balance-related activities. The GVS STIM condition resulted in 5.0% of dreams incorporating arm movements and 10.0% including balance-related activities, with no tactile sensations reported. In the GVS SHAM condition, 12.0% of dreams incorporated arm movements, with 4.0% balance-related activities and no incorporation of tactile sensations. The HS STIM condition showed 21.4% incorporation for arm movements and 7.1% for tactile sensations, with no balance-related activities. The HS SHAM condition had 20.8% of dreams incorporating arm movements, 4.2% including tactile sensations, and no balance-related activities. Overall, the data suggest that EMS and HS stimulation conditions exhibited higher percentages of dreams incorporating arm movements and tactile sensations compared to their respective sham conditions, though these differences were not statistically significant. GVS conditions showed lower incorporation rates for all categories, but no clear conclusions about the relative effectiveness of the different stimulation conditions can be drawn from these results.

A mixed model test was performed to examine the dream incorporation rates for statistically significant differences across conditions, as presented in Table 1. For the Movement of the Arms, there were no significant interaction effects between the conditions (EMS, GVS, HS) and STIM or SHAM.

**Table 1:**
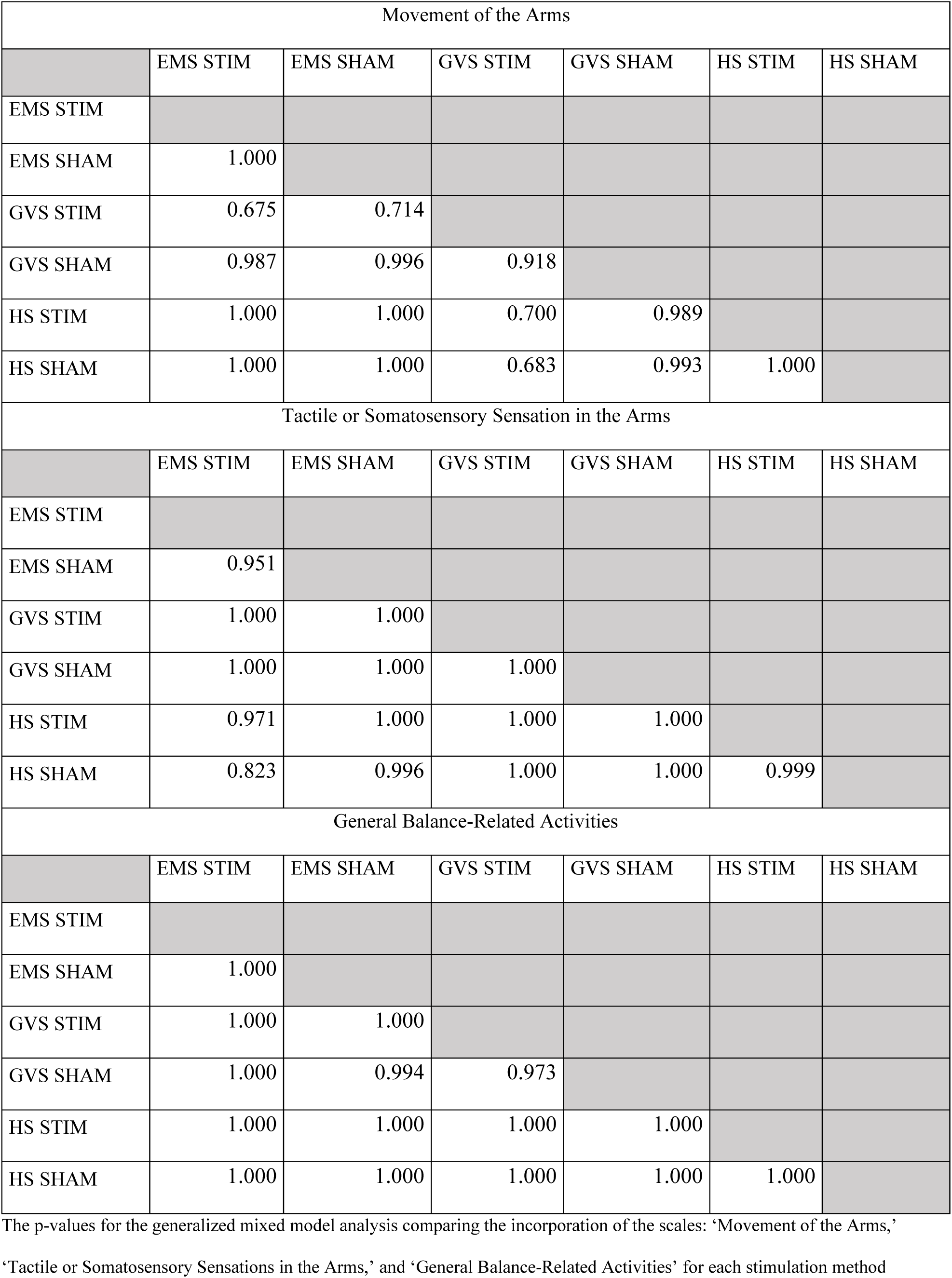
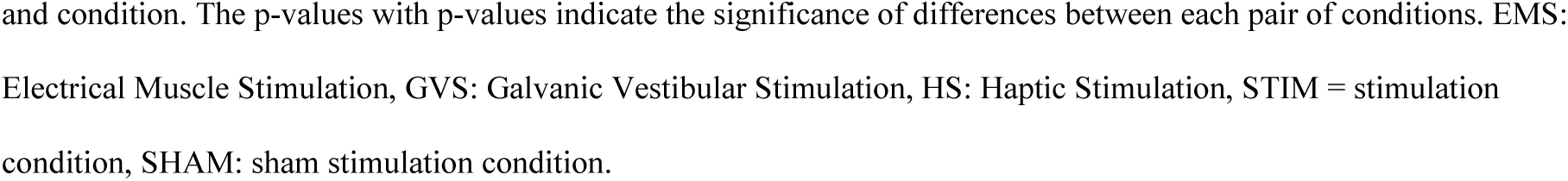
The p-values of the Generalized Linear Mixed Model analysis.

| Movement of the Arms |  |  |  |  |  |  |
| --- | --- | --- | --- | --- | --- | --- |
|  | EMS STIM | EMS SHAM | GVS STIM | GVS SHAM | HS STIM | HS SHAM |
| EMS STIM |  |  |  |  |  |  |
| EMS SHAM | 1.000 |  |  |  |  |  |
| GVS STIM | 0.675 | 0.714 |  |  |  |  |
| GVS SHAM | 0.987 | 0.996 | 0.918 |  |  |  |
| HS STIM | 1.000 | 1.000 | 0.700 | 0.989 |  |  |
| HS SHAM | 1.000 | 1.000 | 0.683 | 0.993 | 1.000 |  |
| Tactile or Somatosensory Sensation in the Arms |  |  |  |  |  |  |
|  | EMS STIM | EMS SHAM | GVS STIM | GVS SHAM | HS STIM | HS SHAM |
| EMS STIM |  |  |  |  |  |  |
| EMS SHAM | 0.951 |  |  |  |  |  |
| GVS STIM | 1.000 | 1.000 |  |  |  |  |
| GVS SHAM | 1.000 | 1.000 | 1.000 |  |  |  |
| HS STIM | 0.971 | 1.000 | 1.000 | 1.000 |  |  |
| HS SHAM | 0.823 | 0.996 | 1.000 | 1.000 | 0.999 |  |
| General Balance-Related Activities |  |  |  |  |  |  |
|  | EMS STIM | EMS SHAM | GVS STIM | GVS SHAM | HS STIM | HS SHAM |
| EMS STIM |  |  |  |  |  |  |
| EMS SHAM | 1.000 |  |  |  |  |  |
| GVS STIM | 1.000 | 1.000 |  |  |  |  |
| GVS SHAM | 1.000 | 0.994 | 0.973 |  |  |  |
| HS STIM | 1.000 | 1.000 | 1.000 | 1.000 |  |  |
| HS SHAM | 1.000 | 1.000 | 1.000 | 1.000 | 1.000 |  |

The p-values for the generalized mixed model analysis comparing the incorporation of the scales: ‘Movement of the Arms,’
‘Tactile or Somatosensory Sensations in the Arms,’ and ‘General Balance-Related Activities’ for each stimulation method
and condition. The p-values with p-values indicate the significance of differences between each pair of conditions. EMS: Electrical Muscle Stimulation, GVS: Galvanic Vestibular Stimulation, HS: Haptic Stimulation, STIM = stimulation condition, SHAM: sham stimulation condition.

None of the pairwise comparisons yielded statistically significant differences for the three categories ‘Movement of the Arms,’ ‘Tactile or Somatosensory Sensations in the Arms,’ and ‘General Balance-Related Activities.’

For contextualization purposes, excerpts of dream reports with clear dream incorporation for each condition are presented in Table 2.

**Table 2:** Dream Incorporation Excerpts.

| Stimulus | Condition | Scale | Dream Report |
| --- | --- | --- | --- |
| EMS | STIM | Movement of the Arms | And I had to stick those adhesive strips on my hand with UV-resistant adhesive tape.' |
| EMS | SHAM | Movement of the Arms | ...it was just a feeling, and I mostly used my right hand to remove, like to move away this white matter. I was just like constantly trying to move it away.' |
| HS | STIM | Movement of the Arms | Then we were just cleaning it. And now I've just been to the fridge and taken out drinks.' |
| HS | SHAM | Movement of the Arms | '...and it was like a box of frozen things. And I started packing them...' |
| GVS | STIM | General Balance-Related Activities | I was on a mission and there we were outside and we were walking up the mountain, but it was shaky.' |
| GVS | SHAM | General Balance-Related Activities | ...and then I fell over. And when I fell over, I fell over into my mother...' |
| HS | STIM | Tactile or Somatosensory Sensations in the Arms | I had my Apple Watch with me and at first I thought it was vibrating.' |
| HS | SHAM | Tactile or Somatosensory Sensations in the Arms | I had to hold the microphone with two hands. I think holding the microphone definitely was something that had a vibration, like an electronic feeling.' |
| EMS | STIM | Tactile or Somatosensory Sensations in the Arms | ...then suddenly I had to get up, and I had something on my wrist.' |
| EMS | SHAM | Tactile or Somatosensory Sensations in the Arms | And there was a couple with a young black labrador. And then I was allowed to pet it.' |

Dream incorporation examples for each stimulation method, condition and the scale this excerpt would qualify
for as Dream Incorporation. Electrical Muscle Stimulation, GVS: Galvanic Vestibular Stimulation, HS: Haptic
Stimulation, STIM = stimulation condition, SHAM: sham stimulation condition.

We calculated the effect size (Cohen’s d) on each scale to explore the magnitude of the difference between groups, independent of sample size resulting in 0.67 in ‘Movement of the Arms’; 0.26 in ‘Tactile of Somatosensory Sensations in the Arms’ and 0.11 in ‘General Balance-Related Activities’. These results indicate that solely based on effect sizes, ‘Movement of the Arms’ could be potentially affected by the different stimulation methods. In contrast, the effect might be less for ‘Tactile of Somatosensory Sensations in the Arms’ and ‘General Balance-Related Activities,’ respectively. Considering the sample size, in our power analysis, with 23 participants, the study was underpowered to detect significant effects, especially for ‘Tactile of Somatosensory Sensations in the Arms’ (power = 6.4%) and ‘General Balance-Related Activities’ (power = 14.1%), where effect sizes were small. Although ‘Movement of the Arms’ showed a moderate to large effect size (power = 78%), the power was slightly below the desired threshold for reliably detecting significance. Our power calculation showed that the study was only capable of detecting relatively large effects. Given the small sample size, smaller effects may not have been detected, potentially contributing to the non-significant results. Therefore, while the lack of significant differences suggests minimal impact, it is possible that limited power influenced the findings.

## Discussion

The effect of three different external stimuli on dream content were investigated on their efficiency to affect dream content.

### Incorporation of Stimuli

#### Galvanic Vestibular Stimulation

We expected GVS to increase the number of balance-related dream references, and the results of the scale ‘General Balance-Related Activities’ did not confirm this hypothesis. In the GVS condition, the STIM condition has 10.0% of DI, and SHAM 4.0% DI. This difference was not significant. Additionally, in 7.7% of the dreams in the EMS SHAM condition, there was a presence of ‘General Balance-Related Activities’, suggesting that the incorporation of ‘General Balance-Related Activities’ was not related to GVS stimulation. Overall, the incorporation rates are lower than in previous work. Leslie and Ogilvie described an incorporation rate of 25% in their study using a hammock, which the results of a GVS strategy in this study do not support. In their study, participants slept in a hammock, which was either stationary (control condition) or rocked at a constant frequency to stimulate the participants’ vestibular system (Leslie & Ogilvie, 1996). In this study, the intrinsic properties of the stimulation method proposed a major challenge as the application of GVS and the resulting perceptions are complex and are dependent on many variables. Firstly, any change in the positioning of the head causes changes in vestibular perception (Khosravi-Hashemi et al., 2019). Participants can experience the feeling of being in a hammock, on a pirate ship, swinging like a pendulum and many other types of vestibular experiences depending on body positioning.

Prior to the experiment, pilot testing was done to explore the position of the head in different sleeping positions and its effect on the perceptions caused by GVS. There was no consistent effect of sleeping positions on the sensations. This shows that the position of the head is an extremely delicate parameter. Even very slight changes in sleeping positions, such as a small shift in head position, could cause different sensations. Only with extremely precise and stable head positioning can this effect be surmounted. However, in a sleep study, this is close to impossible. Not only the spatial position of the head but also the rest of the body influences the subsequent sensation. The signals that are sent to the brain from the vestibular cortex are compared to the tactile and proprioceptive information the rest of the body provides (Proske & Gandevia, 2012). These information streams are then combined and compared, and a conclusion about the body positioning is drawn. This shows us that the position of the entire body, including individual body parts, will influence the vestibular sensation. Finally, together with visual input, a final verdict is given about the true vestibular status of the body (Stein & Stanford, 2008). In wakefulness, the visual input has a large effect on this verdict; however, during sleep and dreaming, there is no external visual input to support this. The effect of externally produced vestibular sensations on visual imagery during dreaming remains an open question. It is argued that visual imagery can produce vestibular or movement sensations (Picard-Deland et al., 2020). External visual input is, in our case, thus not a confounding factor in the production of a vestibular sensation. After testing the bodily positions and different stimulation parameters, as described in the methodology, we decided on a stimulation of 1 Hz with a duration of two seconds. Finally, because of the effects of spatial positioning of the head, it is extremely challenging to create a consistent vestibular effect across participants, even if the stimulation parameters are matched. However, any significant noticeable vestibular effect in the dream might be sufficient for dream incorporation. In the end, the set intensity of the stimulus might have been too low for successful incorporation into the dream. To avoid arousal or premature awakenings, the lowest stimulus intensity was chosen, at which the person would experience any vestibular effect while sitting on the bed during wakefulness. However, it is still unknown whether this threshold is higher or lower during sleep. Also, the length of the stimulus may have been too short in duration for successful incorporation. This length was initially chosen to match the other two stimuli in length, which enables a better comparison between the three. However, the fact that the vestibular stimulation and sensation only lasted for two seconds per stimulus may have suppressed any incorporation effect. The sensation caused by the GVS stimulation during wakefulness resulted in a quick left-right-left-right wiggle from side to side. Perhaps if the stimulus was a more stable, continuous stimulus throughout the entire REM stimulation period, it would have been noticed more and thus resulted in more incorporation. Additionally, the vestibular system is active in almost all the movements we make in our everyday lives (Goldberg et al., 2012); only significantly bigger deviations from “normal” are detected. Thus, the dreamers might have overlooked the small, short vestibular effect. Overall, future research effort is needed to truly understand and tackle the challenging features of GVS.

#### Electrical Muscle Stimulation

In the EMS stimulation condition, there was no increase in dreamed ‘Movements of the Arms’ upon stimulation, showing consistent reporting across all conditions. This could be explained by the fact that EMS stimulation produced a very slight hand movement as a result of a very small current targeting the forearm muscle, however, solely when the arm is unobstructed. As soon as participants caused any obstruction of the hand, may it be due to the pressure of the blanket on the arm, or a certain bodily position, the hand movement could not be produced. In these cases, only the current applied to the skin was felt, without the result of a hand movement. In addition to this, it is methodologically challenging to record whether a hand movement was observed, as we did not have the means to observe what happens under the blankets. The lack of hand movements in real life renders the lack of dreamed arm movement nothing more than expected. Future studies could physically secure the hand in some way, ensuring this hand movement. While this might decrease the quality of sleep in a significant way, it might be worth exploring. For ‘Tactile or Somatosensory Sensations in the Arms’ EMS did not have significantly more dream incorporation upon stimulation (p=0.0378 α=0.0033). This finding will be further discussed in the section: *Electrical Muscle Stimulation and Haptic Stimulation*, as this finding can be viewed in the same light as the results of HS.

#### Haptic Stimulation

HS stimulation did not evoke more incorporation when compared to sham stimulation for ‘Movement of the Arms’ (p=0.067, α=0.0033) and ‘Tactile or Somatosensory Sensations in the Arms’ (p=0.067, α=0.0033). The use of HS as a stimulation method in modern times, particularly with smartwatches, introduces two potential effects: sensitization and desensitization. Participants may be sensitized, leading to heightened responsiveness and arousal due to associating wrist vibrations with an alarm or a message notification. Conversely, they may be desensitized, perceiving the stimulation as familiar and thus less likely to incorporate it into their dreams. Researchers should consider these factors when using HS as a method during sleep experiments. To overcome these effects, future studies can apply HS to a different extremity. A vibrational stimulus on the ankle of participants was applied to induce lucid dreaming (Paul et al., 2014). They achieved higher lucid dream induction and incorporation with stimulation of the ankle compared to the wrist. The participants were able to correctly guess the body part that was stimulated most of the time. However, this study did not use any control condition. It is thus hard to say what the true incorporation rates are without any sham stimulation.

#### Electrical Muscle Stimulation and Haptic Stimulation

Because the actual hand movement was occasionally obstructed in the EMS condition, EMS and HS seemed to be of quite a comparable nature. EMS and HS overlap in their quality of sensation, as both stimuli were attached to a part of the forearm. Both can have various qualities such as pressure, touch, or vibration. Each of these qualities is recognized by different receptors such as the Merkel Cells, Meissner’s Corpuscles, Pacinian Corpuscles, or free nerve endings (Bartels & Bartels, 2004; Menche et al., 2023). The activation of several receptor types could result in a lot of afferent information to the brain and may be an influence on incorporation. Furthermore, depending on the place of stimulation, the density of these receptors can be very high (e.g. hand) or less high (e.g. inner forearm in our case. This could be important, since a too “receptor-dense” location may lead to more awakenings instead of incorporations (Paul et al., 2014). This framework applies to both HS and EMS, however, one significant difference remains. EMS stimulates the muscle directly and could be considered more invasive than HS, which only stimulates the surface of the skin. However, our results do not support this hypothesis for the scale ‘Tactile or Somatosensory Sensations in the Arms’. The fact that the SHAM condition evoked similar incorporation for ‘Movement of the Arms’ in both the HS and EMS speaks for the significant role of the ‘theatre-effect’ of the experiment itself on dream incorporation. Solely being in the lab with the researcher, while wearing strange devices on the body might have a strong effect on dream content regardless of whether an actual stimulation was performed. In addition, the fact that the sensation of having electrodes on the skin (with or without electrical or haptic stimulation) could be incorporated in the dream. It is a tactile stimulation by itself, as there are electrodes on the body.

Regarding the effectiveness of any chosen stimulation, Schredl (2018) proposes different factors: The stimuli being physically close to oneself while sleeping, e.g. being tactile as opposed to audio and/or visual stimuli, could be more important and hence processed differently. They may also carry a bigger immediate threat, while still maintaining the balance of not waking the person (Schredl, 2018; Solomonova & Carr, 2019). Another potential confounding factor could be that a REM awakening with stimulation could have an effect on the following SHAM condition. Even though there is no stimulation in that REM phase, the memory of the stimulation that occurred in the previous REM phase might cause incorporation.

### Limitations

#### Data Acquisition

Beyond limitations that are specific to the stimulation methods, the study design contained more general limitations. There was no blinding of the participants as to which kind of stimulus they would receive on which night. They were, however, blind to the STIM/SHAM order. The decision against fully blinding the participants was made for two reasons. Firstly, sleeping with all three devices attached during all four nights could’ve possibly worsened the sleep quality of the participants as sleeping in a strange environment can already limit sleep quality (Edinger et al., 1997). In contrast, the lack of blindness to the type of stimulus might not be regarded as a limitation, as the goal of the study was to have a high incorporation of the stimuli. Priming the participants pre-sleep to pay attention to a certain stimulus only aids this goal and by using a sham control condition, the incorporation rates are still meaningful. A more rigorous approach would be to conduct a double-blind experiment, in which the participants, as well as the researchers, are both unaware of the condition. In our study design, this was discussed; however, due to specific EEG artifacts caused by both GVS and EMS, but not HS, double blindness was, in our case, impossible. Another point of improvement would be to test the stimulation parameters in the adaptation night during sleep, to establish the correct intensities. The stimulus intensity should have been increased until arousal and this way, the most accurate intensity would have been selected. Since this was not done, it may have reduced the number of successful incorporations. In studies on dream incorporation and lucid dream induction, assessing wake thresholds is crucial for optimizing data acquisition and outcomes, but it presents significant challenges. Researchers are currently developing individualized arousal detection algorithms that adjust stimulus intensity based on real-time online PSG data. This advancement aims to significantly mitigate the wake threshold issue (Pavlou, 2024).

The dream report collection after the REM awakenings was relatively simple with an open question and a direct question targeting the stimulus specifically. This was decided as any additional questionnaires increases the length of the awakening and the arousal level of the participants, making it more challenging to fall asleep quickly afterwards. In addition to open dream reports, Noreika et al. (2020) collected very specific questionnaire data, which the participants wrote down right after being awakened. They found differences when comparing the free dream reports to the questionnaire data, regarding the effects of tDCS on repetitive movements. They speculated that some bodily experiences might only be reported when specifically asked for (Noreika et al., 2020). Having a questionnaire that collects a variety of dimensions may be better suited to detect the incorporation of stimuli; thus, future studies would benefit from adding more targeted self-reported scales.

#### Dream Content Analysis

As mentioned in the methods section of this study, dream reports that included any indication of the participant being woken up from stimulation were excluded. In addition to this, the PSG data was assessed on the presence of wakefulness during stimulation. During this process, a conservative approach was taken, and if the situation was unclear, the dream reports were excluded. This might have increased the number of false negatives. Additionally, in dream content analysis, there is usually a differentiation between so-called direct and indirect incorporation. Direct incorporation rates are rather easy to score. When applying the stimulus while sleeping (e.g. vibration on a finger), direct incorporation would be the finger vibrating in the dream (Paul et al., 2014). While direct incorporation is relatively straightforward, uncertainty arises when assessing indirect incorporation. When applied, vibration can affect the dream content in various ways. One example of this could be the whole body vibrating, but it could also be conveyed in an earthquake, or even a sudden change of scenery. These implicit potential consequences make indirect incorporation much more challenging to objectively score. Hence, the final scoring is very dependent on the set scoring rules and formulation of the dream content scales. Additionally, some dream content can be scored as incorporation for multiple scales. This is shown in the excerpt on dream incorporation in table 2. The participant mentions ‘…and there was a couple with a young black Labrador. And then I was allowed to pet it.’ In our investigation, we wanted to look at dreamed arm movements, as well as tactile sensations on the arm. Petting a dog would be scored as incorporation for both scales, as petting a dog is a movement of the arm and simultaneously evokes a tactile sensation. Thus, the results from ‘Movement of the Arms’ and ‘Tactile or Somatosensory Sensations in the Arms’ will contain overlapping data. Another interesting factor that we did not introduce into our methodology is the fact that arousal thresholds are different in tonic versus phasic REM sleep (Ermis et al., 2010). We looked at dream incorporation in all REM stages thus making no distinction between the two. In future research, tonic and phasic REM sleep can be separated, and based on this, the stimulation thresholds can be altered accordingly, optimizing the potential for dream incorporation while avoiding wakefulness.

### Contributions to the Study of Dreams, Consciousness, and Neuroscience

Understanding the interplay between internal and external factors in dream formation reveals insights into the complexities of consciousness during sleep. Aligned with the concept of ‘hybrid states of consciousness,’ where waking and dreaming boundaries blur (Revonsuo, 2006), dream incorporation raises questions about how external stimuli impact memory consolidation. This work suggests that factors like timing, stimulus intensity, and emotional significance shape memory outcomes during sleep, emphasizing the intricate and context-dependent nature of dream incorporation’s effects on memory function (Cairney et al., 2016; Diekelmann & Born, 2010). It is unknown how the three stimuli are truly perceived in the dream world and how they affect the dream body. Lucid dreaming could provide a solution to this problem. Lucid dreaming is an exceptional state of consciousness defined as having wake-like awareness during a dream that enables the potential to have a degree of control over the dream. This unique brain state has earned significant interest due to its potential applications in learning, creativity, mental well-being, and motor learning (Erlacher & Schredl, 2010; Peters et al., 2023). By presenting the stimulus during a lucid dream, participants can be instructed to notice the stimulus closely and describe the sensation in detail using a verbal dream report. Taking this a step further would be the use of two-way communication. In 2021 four multi-site studies researched the feasibility of live two-way communication during a lucid dream using four different approaches (Konkoly et al., 2021). This shows us that besides a detailed dream report, live communication about the stimulation is possible while it is happening. At this point in time, the possibilities of communication are limited to more simple methods, like time stamping, counting, Morse code, and yes/no answers. However, the future of dream communication studies is filled with creative and innovative ideas on how to elaborate these methods further. Furthermore, by investigating the incorporation of external somatosensory stimulation into dream content, this study lays the foundation for potential applications in lucid dreaming induction. The identification of specific stimuli that act as cues for triggering lucidity during dreams via Targeted Lucidity Reactivation (TLR), could have far-reaching implications for cognitive enhancement, psychological therapy, and self-exploration (Carr et al., 2020). In addition, the integration of external stimuli into dreams has parallels with virtual reality experiences, where external sensory input can create immersive, seemingly real environments. Exploring the similarities and differences between dream incorporation and virtual reality can deepen our understanding of both phenomena, how to combine them, and finally, how to optimize lucid dream induction using virtual reality (Gott et al., 2020). This study goes beyond dreams, highlighting significant medical implications, particularly in nightmare therapy and mental health. By exploring how external stimuli affect dream content, it opens doors for therapeutic interventions like Targeted Memory Reactivation and Imagery Rehearsal Therapy, providing potential relief for individuals with distressing dreams, nightmares, or PTSD. Investigating the controlled application of somatosensory stimuli during sleep could maximize the efficacy of these therapeutic measures.

## Conclusion

The investigation of incorporating external somatosensory stimulation into dream content represents a fundamental contribution to various scientific fields. By investigating the relationship between external stimuli and dream formation, this research strengthens our understanding of the fundamentals of dreaming, memory, consciousness science, nightmare treatment, and the potential for lucid dreaming induction.

Our study’s meticulous design, including the comparison of multiple stimuli and a thorough control condition, addresses many of the complexities often overlooked in simpler experimental frameworks. Although our findings indicate that these methods did not produce strong effects on dream incorporation, this outcome highlights the inherent challenges of designing rigorous dream research studies. The outcome of our power analysis suggests that the methods employed are unlikely to induce strong effects on dream incorporation, particularly in the context of lucid dream induction. This finding has practical implications, as methods that require strong effects to be useful, such as for consumer applications, may not find these stimuli effective. Nevertheless, we cannot exclude the possibility that the stimuli might exert weak or medium effects, which could still be relevant in combination with other strategies or for certain individuals, and the comprehensive approach provides valuable insights into the methodological challenges and nuances of dream incorporation research. Refining stimulation methods and exploring individualized applications could enhance the effectiveness of dream incorporation techniques. By setting a high standard for experimental design, our study makes a valuable contribution to dream research and related fields as it provides a solid foundation for future work aimed at improving methods for studying dreams.

## Acknowledgments

I would like to extend my sincere thanks to the individuals who contributed to the completion of this manuscript by providing valuable proofreading and insightful feedback, which greatly improved the quality of the work. I am also deeply grateful to the students at the Institute of Sports Science at the University of Bern, Switzerland, who helped collect and analyze the data, including their dedication to overnight recordings and assistance in working out the dream data. Their efforts were essential to the success of this research.

## Data availability statement

The data supporting the findings of this study will be made available in the BORIS repository of the University of Bern.

